# Predicting Conscious Perception from Pupil’s Aperture Size Using Machine Learning Techniques

**DOI:** 10.64898/2026.08.26.747446

**Authors:** Pragya Pandey, Sayali Raveendra Pethe, Indrajeet Indrajeet, Supriya Ray

**Affiliations:** Integrative Neuroscience and Cognition Centre, CNRS UMR 8002, Université Paris Cité, France; Centre of Behavioural and Cognitive Science, University of Allahabad, India; Department of Psychology & Cognitive Sciences, Ashoka University, Plot #2, Rajiv Gandhi Education City, Sonipat 132029, Haryana, India

**Keywords:** Perception, Decision, Attention, Pupillometry, Machine Learning

## Abstract

**Introduction:** Decision making for selecting an object or a course of action from possible alternatives largely depends on our perceptual ability modulated by attention. When multiple stimuli appear close together in time, processing one stimulus can temporarily impair the processing of another due to temporal limitations of attention. Observers frequently fail to detect the second target (T2) presented within a few hundred milliseconds after the first target (T1) in a stream of stimuli, which is commonly known as ‘attentional blink’ (AB). Existing theories attribute this perceptual lapse to T1 processing, distractor interference, or transient attentional gating; however, the computations underlying suppressive mechanism remains unresolved. We investigated whether pupil-size could reveal the underlying mechanisms of AB and predict conscious perception on a trial-by-trial basis.

**Methods:** Pupil diameter and gaze locations were recorded using an infrared eye tracker. Machine learning techniques were used to classify trials when T2 was detected versus when it was not, after correct identification of T1, during an AB task from the pupil dynamics, which also yielded attentional episode (AE) associated with each element in the stream of visual stimuli when deconvolved.

**Results:** Cross-validating classifiers achieved near-perfect accuracy not only in distinguishing but also predicting perceptual outcomes on a single-trial basis. AEs exhibited greater power when T2 was detected than when it was missed; the differential power in AEs on a logarithmic scale was highly synced with the differential pupil size.

**Conclusions:** Collectively, these findings establish a framework for predicting attention-driven perceptual outcomes from pupil-dynamics at finer time-scale.

## Introduction

The temporal limits of conscious processing of visual information are commonly examined using the Rapid Serial Visual Presentation (RSVP) task in which two visual targets are presented in quick succession within a rapid stream of distractors. Human participants often fail to detect the second target (T2) when it appears 200–500 ms after the first target (T1), a phenomenon known as the attentional blink (AB) [1,2].

Although the dynamics of pupil size averaged across several trials differed between trials in which human participants reported T2 (i.e., no-blink trials) and trials in which T2 escaped conscious perception (i.e., blink trials), the classification of individual blink and no-blink trials based on pupillary responses has not yet been explored [3]. Using ‘Support Vector Machine (SVM)’ and ‘Logistic Regression (LR)’ classifiers in MATLAB^®^ statistics and machine learning toolbox, we were able to predict perceptual outcomes from pupil dynamics with almost absolute certainty [4]. SVM identified an optimal decision boundary that maximized separation between two classes, whereas LR estimated class membership probabilities by applying a sigmoid function to a linear combination of input features.

The pupil size is considered an indirect marker of locus coeruleus (LC) activity [5]. The LC is a small bilateral nucleus located in the dorsal pons of the brainstem that releases norepinephrine (NE) throughout the cortex. The LC-NE neuromodulatory system is the brain’s primary source of cortical norepinephrine, crucial for regulating arousal and attention [6–8]. Nieuwenhuis and colleagues proposed that the AB arises from target-driven attentional enhancement mediated by the LC–NE system; the detection of T1 triggers a transient burst of NE that temporarily enhances attentional processing [9]. However, this enhancement is followed by a refractory period, consequently, when T2 appears during this interval, it fails to receive sufficient attentional enhancement, increasing the likelihood that it will escape conscious report. These alternating periods of attentional enhancement are termed attentional episodes (AE) [10,11].

By deconvolving pupillary responses recorded during the RSVP task using a pupillary impulse response function, we reconstructed the timing and magnitude of transient underlying attentional events (AEs). Unlike other contemporary theories that posit relatively longer AEs triggered only by T1 and T2 [10,12,13], the deconvolution method naturally yielded us AEs of shorter duration triggered by every item in the RSVP, which were represented by brief oscillatory signals. Taken together our study shows that relative strength in the AEs determine the perceptual outcome, which can be predicted from the pupil dynamics on a trial-by-trial basis.

## Materials and Methods

Twenty healthy adults with normal or corrected-to-normal vision participated in the RSVP detection task (7 females, 13 males; mean age ± SD: 21.8 ± 2.14 years). Due to droopy eyes and / or excessive blinking, three subjects did not contribute data in the final analyses. A written informed consent was obtained from every participant after explaining the protocol, and all data were anonymized to ensure confidentiality. Instructions were provided verbally and in writing in either Hindi or English, according to participant preference.

Participants completed 50–100 practice trials before recording sessions and received fixed monetary compensation with performance-based incentives. The study adhered to ICMR Ethical Guidelines (2006), consistent with the Declaration of Helsinki, and was approved by the Institutional Ethics Review Board of the University of Allahabad.

The study employed a variant of rapid serial visual presentation (RSVP) task, commonly known as the ‘attentional blink’ task (1). This task was used earlier elsewhere [14]. Stimuli were presented on a 1024 × 768, 60 Hz LCD monitor, while gaze position and pupil diameter were recorded monocularly at 60 Hz using an ASL D6 video based infra-red eye tracker (Applied Science Laboratories, MA, USA) (1). Programs written in E- Basic language of E-Prime 1.2 software (Psychology Software Tools Inc., USA) presented the stimuli, monitored the gaze positions, time-stamped the task related events, and provided the feedback of performance. Each trial began with a central grey fixation spot (∼0.4° × 0.4°) on a black background, accompanied by two peripheral green squares positioned 13° from fixation in both hemifields. Stimulus presentation commenced only after the participants had maintained gaze within a ∼2° × 2° fixation window for ∼500 ms. The RSVP stream consisted of uppercase English letters presented centrally within ∼0.9° × 0.9° visual angle. Letters were randomly selected without replacement from the alphabet excluding I, O, Q, S, and Z to avoid visual similarity with numerals. Each item was displayed for ∼34 ms followed by a blank interval of ∼51 ms, with an additional temporal jitter of 0–2 refresh durations, yielding an average ISI of ∼85 ms (12 items/s). All letters were grey and equiluminant. After 4–8 letters, a white numeral (T1; randomly selected from 2, 3, 4, or 7) appeared and participants classified it as even or odd. In 50% of trials, the second target (T2), a grey letter “X”, appeared after T1 at lags 1, 2, 3, 4, 6, or 8, where lag-n indicates (n – 1) distractors appeared between T1 and T2. The RSVP stream continued for 2–5 additional letters after T2. Participants were instructed to maintain fixation throughout; online eye tracking aborted trials with fixation breaks. The luminance of fixation, peripheral squares, T1, letters (including T2), and background was 3.45, 2.21, 18.5, 1.31, and 0.24 cd/m², respectively.

Each participant completed 240 trials of the RSVP task, in which they classified the numeral target (T1) as odd or even and reported the presence or absence of the letter ‘X’ (T2). Following the RSVP stream, participants first responded to the T1 classification task by pressing the left mouse button for an even numeral (2 or 4) and the right mouse button for an odd numeral (3 or 7). They then responded to the T2 detection task by pressing the left mouse button if the letter ‘X’ was detected and the right mouse button otherwise. A maximum response window of 10 s was provided for each decision. After each response, visual feedback (“Correct” or “Incorrect”) corresponding to both T1 classification and T2 detection was presented on the screen.

The in-house programs written in MATLAB^®^ (The Mathworks Inc., USA) were used to analyze the pupil and eye-position data offline. All statistical calculations were performed using either MATLAB^®^ Statistical Toolbox (The Mathworks Inc., USA) or SigmaStat (SYSTAT Inc.,USA). Trials containing missing values, discontinuities, or extreme task-irrelevant pupillary fluctuations—likely caused by blinks or partial eye occlusions—were identified and excluded. Trials showing a maximum–minimum pupil difference greater than 50 units were filtered using a Hampel filter (window length = 5) and median interpolation (2). Trials still exceeding this threshold after filtering were removed. To reduce the influence of noise and artifacts, pupil size was normalized using a baseline corrected divisive normalization, with the average pupil diameter during a baseline period (∼400 ms following gaze fixation) serving as the denominator (i.e., 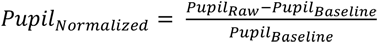). We aligned normalized pupillary response either at the time of RSVP onset or the first target (T1) onset. Further, a discrete Fourier transform was performed on the pupil data from each trial to generate the signal power spectrum. Finally, the conventional interquartile range (IQR) method was used to remove outliers from the power spectrum. To understand pupillary dynamics in the RSVP task, trials were pooled across 17 subjects. We also measured pupil size and pupillary light reflex of nine people in response to a flash of light using a handheld pupilometer (IDMED NeuroLight, France). The device uses infrared imaging and a controlled light stimulus to record pupillary dynamics with high precision.

We used machine learning algorithm to evaluate how reliably pupil dynamics can predict the perceptual outcome in a RSVP task. To this end, first, the pupil dynamics in each trial was approximated by fitting a pair of Gaussian functions, 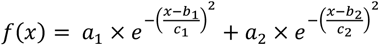 using MATLAB^®^ curve fitting and optimization toolboxes. Next, we extracted features of the pupil dynamics from fitting coefficients *a1*, *b1*, *c1*, *a2*, *b2*, and *c2*. Subsequently, we conducted principal component analysis (PCA) on z-scores of four features: *a*1, *a*2, *b*1/*c*1 and *b*2/*c*2. The ratio ***b***/***c*** represented the normalized center position of the peak relative to its width, i.e., how precisely the location of the Gaussian peak can be estimated in the presence of noise. A high ***b***/***c*** ratio indicates a peak that is far from the origin relative to its own spread. The PCA is a dimensionality reduction technique that transforms large datasets into a smaller set of meaningful factors while preserving most of the original information. The first principal component (PC1) captures the greatest variance in the data, and the second principal component (PC2) captures the next highest variance. The Kaiser-Meyer-Olkin (KMO) index, which measures the sampling adequacy was found 0.6071 for blink trials, and 0.5178 for no-blink trials, with all of the diagonal elements of the anti-image correlation matrix greater than 0.5 (max: 0.6663, min: 0.5002 for blink trials; max: 0.7769, min: 0.5109 for no-blink trials) indicating the number of trials was just adequate for PCA.

We used a linear-kernel support vector machine (SVM) and a logistic-regression (LR) classifier for binary classification of the clusters of data-points in the two-dimensional PCA feature space. The 10-fold cross-validation resampling technique was used for both models to obtain decision boundaries. A K-fold cross-validation resampling technique involves partitioning the dataset into K mutually exclusive subsets (or folds) of equal size; the model is iteratively trained on (K – 1) folds and validated against the remaining fold. This process is repeated K times, such that each fold serves as the independent test set exactly once. By aggregating the performance metrics across all K iterations, the method provides a stable estimate of the model’s efficacy while significantly reducing the variance and bias associated with a single train-test split.

## Results

A total of 17 young healthy participants, who performed a RSVP task (Figure 1A), contributed data for analyses. Trials were grouped by SOA (stimulus onset asynchrony, or the delay between onsets of targets): short (0–183 ms), medium (184–349 ms), and long (350–782 ms). Across all three SOA groups, in some trials participants correctly judged the parity of T1 and successfully detected T2 (“no-blink” trials), while in others they correctly identified the parity of T1 but missed T2 (“blink” trials). Figure 1B shows T2 detection in the RSVP task. Mean (± SEM) success rates were 57.2% (±3.2), 84.6% (±1.9), and 85.2% (±2.1) for short, medium, and long SOAs, respectively. There was a significant main effect of SOA on T2 detection, indicating attentional blink (AB). The Kruskal-Wallis H test indicated that there is a significant difference in the T2 detection performance between different groups of SOAs [χ^2^ (2) = 23.72, p < .001, η^2^ = 0.45], with a mean rank score of 11.71 for short-SOA, 32.18 for medium-SOA, 34.12 for long-SOA. The Post-Hoc Mann Whitney U test using a Sidak corrected alpha of 0.017 indicated that the mean ranks of the short-SOA group is significantly different from medium-SOA and long-SOA groups.

**Figure 1:**
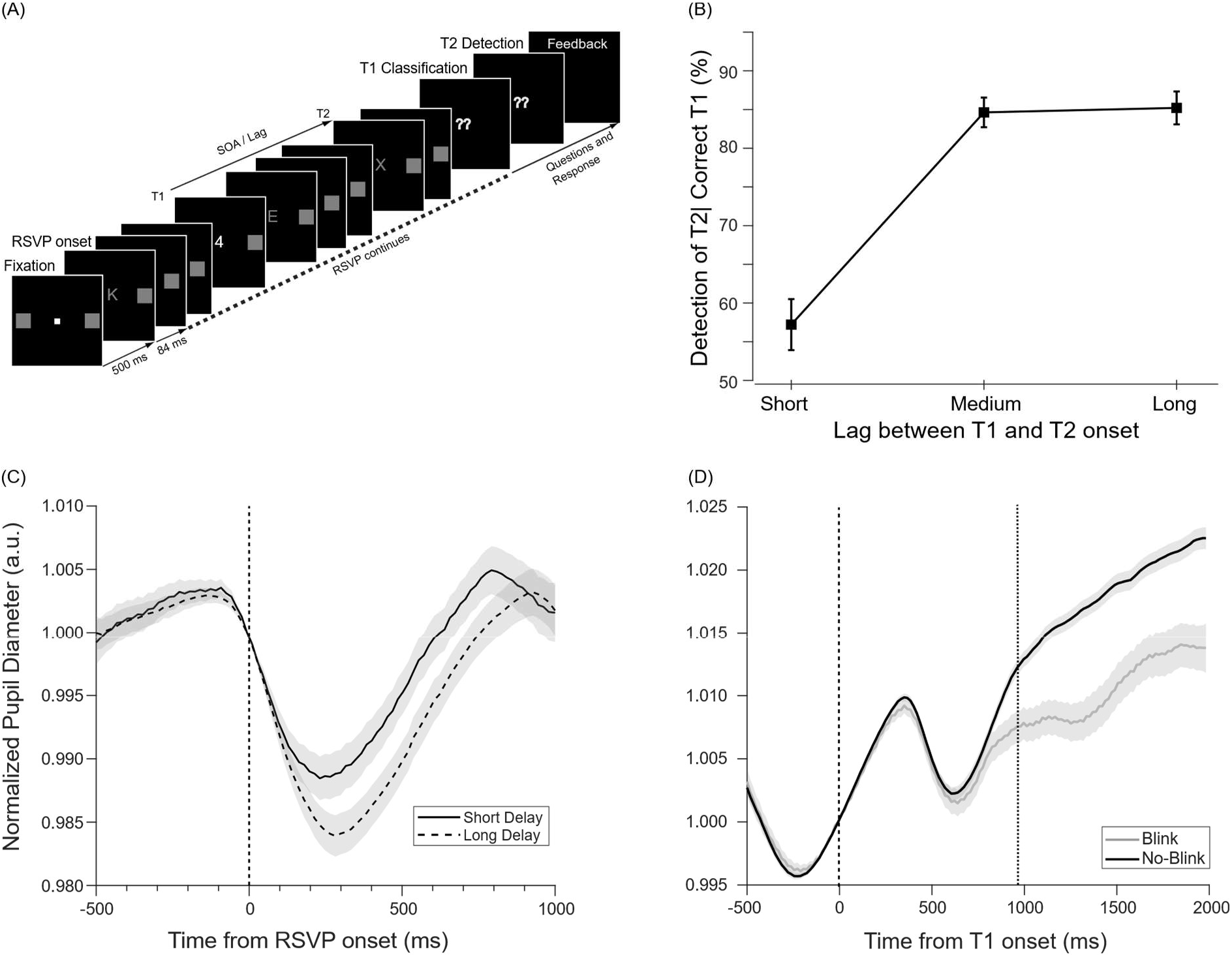
**(A)** In a ‘Rapid Serial Visual Presentation’ (RSVP) task, two targets – T1 (a numeral painted in white) and T2 (letter ‘X’) – were embedded within a stream of English alphabet letters, all painted in grey. The T2 appeared only in randomly selected half of the total number of trials. Participants were asked to identify the parity of T1 (i.e., ‘odd’ or ‘even’) and report T2 (i.e., present or absent). **(B)** Participants exhibited ‘attentional blink’ (i.e., failed to report T2) when T2 appeared soon after T1. **(C)** The first phase of pupil dilation after constriction caused by RSVP onset was delayed when T1 appeared later in the RSVP stream. **(D)** Differential pupil dynamics for blink (grey: T2 missed) versus no-blink (black: T2 detected) trials. In both types of trials, T1was classified correctly. Divergence in pupil size occurred in the second phase of dilation. The vertical dotted line indicates the time when the pupil dynamics statistically differed in two types of trials, starting at 979 ms after T1 onset (black dashed line).

We analyzed pupil dynamics observed during the performance of the task to test whether the modulations in the pupillary aperture reflected subjects’ ability to perceive targets within a rapid stream of distractors. We pooled trials from all participants wherein only T1 appeared that was correctly identified. Trials were then split into two groups based on mean (± SEM) T1 onset delay relative to RSVP: early [413.79 ± 5.83 ms] and late [604.88 ± 5.42 ms]. The delay differed significantly between groups [p < 0.001, t = -23.61, Cohen’s *d* = 3.22]. Baseline-corrected, normalized pupil diameter was aligned to RSVP onset. The mean (± SEM) normalized pupil diameter (Early: 0.991± 0.0014, Late: 0.988 ± 0.0012) and the average (± SEM) height of the peak pupil size (Early: 1.004± 0.0008, Late: 1.002 ± 0.0006) within the first 1 sec after RSVP onset did not differ between groups [mean pupil diameter: p = 0.079, t = 1.77; peak pupil size: p = 0.089, t = 1.71], but the peak of the pupillary dilation averaged across the trials was delayed by 136 ms when T1 appeared late. This suggests that post-constriction dilation following RSVP onset was linked to T1 timing (Fig. 1C).

Subsequently, we segregated trials pooled from all participants into two groups: blink and no-blink. Figure 1D shows differences in pupil dynamics between no-blink (black) and blink (grey) trials. The average (± SEM) normalized pupil diameter is plotted from 0.5 s before to 2 s after T1 onset for each condition. We defined the time of differential pupil dynamics as the point when pupil diameter reliably distinguishes no-blink (black) from blink (grey) trials for at least ∼200 ms (12 consecutive samples). Pupil diameter was significantly larger in no-blink trials (P < 0.05), starting at 979 ms after T1 onset. A clear distinction in pupil dynamics between blink and no-blink trials encouraged us to investigate whether the ability to detect T2 in a rapid stream, with correct T1 identification, could be predicted from trial-by-trial fluctuations in pupil diameter.

We extracted features of the pupil dynamics in each trial by fitting a pair of Gaussian functions, 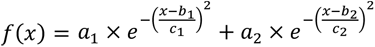, where the fitting parameter ***a*** is the amplitude or the height of the peak of the curve, ***b*** is the mean or position of the center of the peak on the x-axis, and ***c*** is the peak’s width and proportional to the standard deviation. The best fits are overlaid on normalized pupil dynamics in two exemplary trials, one blink and the other no-blink (Figure2A). The pair of Gaussian functions accounted for the biphasic dilation related to T1 and T2 perception. The average (± SEM) goodness of fit (R^2^) across 273 blink trials was 0.687 ± 0.016, and that across 1364 no-blink trials was 0.771 ± 0.006. From the best fit pupil dynamics in each trial, we calculated ***b***/***c***; the ratio represented the normalized center position of the peak relative to its width, i.e., how precisely the location of the Gaussian peak can be estimated in the presence of noise. A high ***b***/***c*** ratio indicates a peak that is far from the origin relative to its own spread. Thus, the dynamics of the pupillary aperture in each trial, both blink trial and no-blink trial, was characterized by four features: *a*1, *a*2, *b*1/*c*1 and *b*2/*c*2.

Next, we centered the data by calculating z-scores of the corresponding feature, and conducted principal component analysis (PCA) on all z-scored features of pupil dynamics in blink trials and no-blink trials using Singular Value Decomposition (SVD) algorithm. Bartlett’s Test for equality of variances showed that the assumption of homoscedasticity was met across four feature dimensions for both blink and no-blink trials [*χ^2^*(3) >> 0.05]. The outliers were removed using the Generalized Extreme Studentized Deviate (GESD) algorithm, and the remaining datapoints were plotted in the 2-dimensional PCA projected feature-space (Figure2B). Merely 7.69 % data points in blink trials and 7.99% data points in no-blink trials were identified as outliers. In blink trials (magenta dots), the first principal component (i.e., PC1) accounted for 54% variance, and the second principal component (i.e., PC2) accounted for 25% variance. In no-blink trials (blue dots), PC1 accounted for 48% variance, and PC2 accounted for 25% variance. Statistically significant differences were found between the two clusters [One-way MANOVA: Wilks’Ʌ (Lambda) = 0.151, F(2, 1504) = 4230.594, p < .001, η^2^= 0.849]. To visualize group clustering, a 95% chi-squared (χ2) confidence ellipse (CE) was generated and overlaid on the PC-space for each set of datapoints originated from blink and no-blink trials. This CE defines a region where roughly 95% of data points to fall, assuming a bivariate normal distribution. The centre of the CE corresponding to blink trials was at (-0.1434, -0.1427) marked by a black ‘X’, and at (-0.1749, -0.0396) marked by a grey ‘X’ for no-blink trials are also shown in Figure2B. The Mahalanobis distance between the two clusters’ centroids was large (D = 40.9), indicating minimal overlap, if there was at all any.

Subsequently, we trained a linear-kernel support vector machine (SVM) and a logistic regression (LR) classifier using 10-fold cross-validation and a default decision threshold of 0.5 for binary classification. To account for class imbalance, sample weights proportional to the square root of the class ratio were applied during training. Thus, the cost of misclassification of no-blink as blink was set almost twice the cost of misclassification of blink as no-blink. The resulting decision boundary clearly separated the data points in the two-dimensional PCA feature space, into distinct clusters with minimal overlap between them (Figure 2B). For the SVM model, a total of 251 support vectors (black circles) defined the margins; the model achieved 95.4% balanced accuracy. The LR model achieved a balanced accuracy of 92.3 %. The LR decision boundary (grey line) is shown in Figure 2B with SVM decision boundary (black line) for comparison.

**Figure 2:**
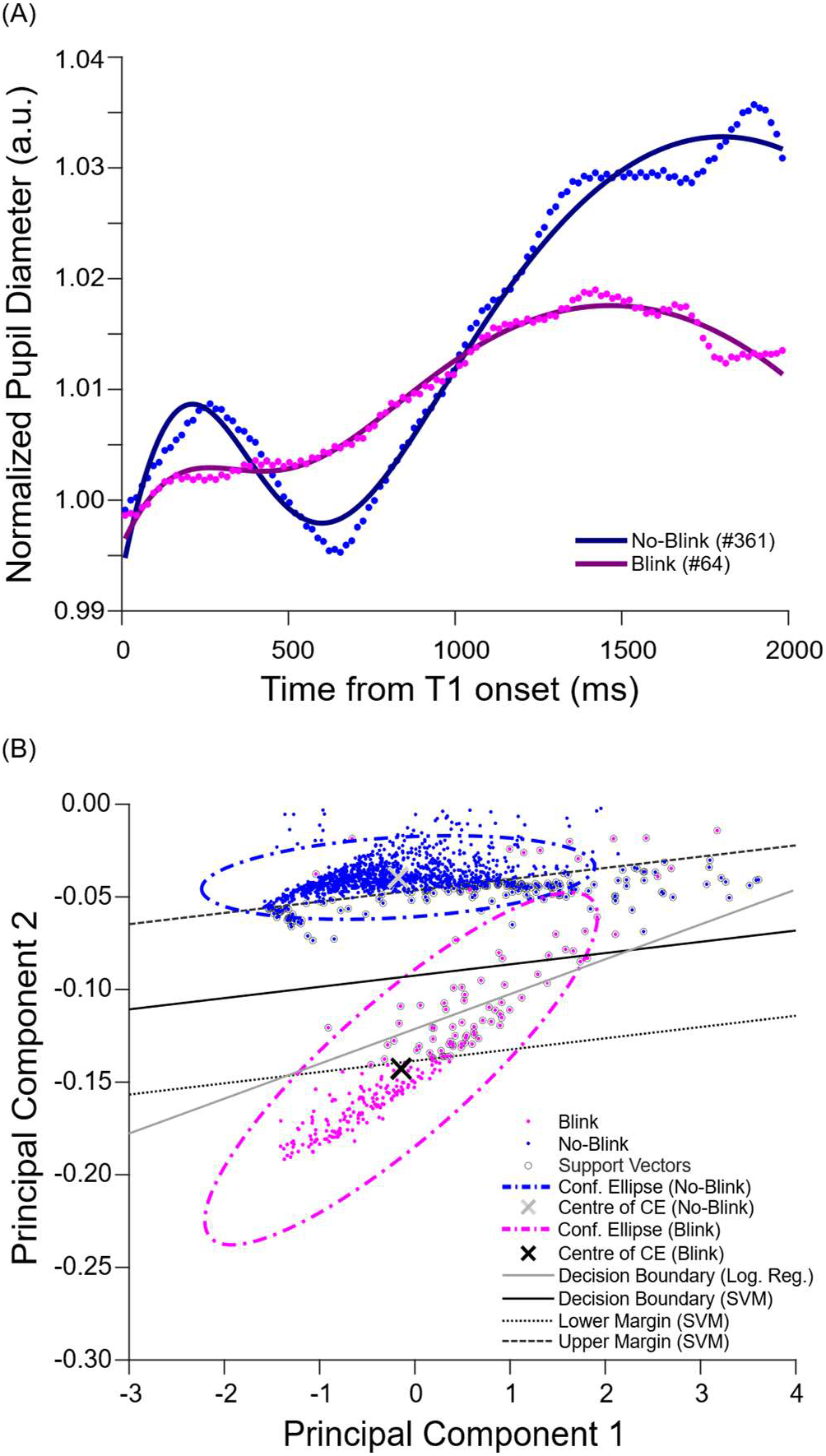
**(A)** The best fit pupil dynamics during a representative blink (purple line) trial and a no-blink (navy blue line) trial. Fitted lines are overlaid on fluctuations in pupillary aperture (blink: magenta, no-blink: blue). **(B)** Plot of pupil-dynamics features on principal component space. Each point was originated from either blink (magenta circles) or no-blink (blue circles) trial. Clusters corresponding to perceptual outcomes in the RSVP task were clearly separable within the two-dimensional PCA-projected feature space of pupil dynamics. The 95% confidence ellipse (dash-dot) for both types of trials (blink: magenta, no-blink: blue) are overlaid on the corresponding clusters. Centroids of confidence ellipses are shown as ‘X’ (blink: black, no-blink: grey). The decision boundary (black line) along with the upper margin (black dashed lines) and lower margin (black dotted lines) obtained from the support vector machine (SVM) classifier are plotted. The decision boundary obtained from the logistic regression classifier is also plotted as a grey line. Support vectors are shown by black circles.

We calculated ‘accuracy’ (i.e., the proportion of correctly classified samples), ‘sensitivity’ or ‘true positive rate’, and ‘specificity’ or ‘true negative rate’. We also calculated F1 score, which is the harmonic mean of precision (i.e., of all the instances the model predicted as ‘blink’, how many were actually ‘blink’?) and sensitivity (i.e., of all the actual ‘blink’ instances that exist, how many did the model correctly identify?).

Because no-blink trials were five times more than blink trials, mere ‘accuracy’ could be misleading, but the F1 score provided a more balanced view. The classification and prediction performance of the SVM and LR were evaluated using Receiver Operating Characteristic (ROC) analysis and a Confusion Matrix respectively. The area under the ROC curve (AUC) quantifies a model’s ability to discriminate between classes across all possible decision thresholds. A confusion matrix evaluates how well a classifier predicts class labels by comparing the predicted outcomes with the true outcomes. These analyses revealed a robust difference between pupil dynamics in two perceptual states (i.e., blink or missing T2, and no-blink or correct detection of T2) in the two-dimensional PC space. Performance of SVM model and LR model are contrasted in Table 1.

**Table 1.**
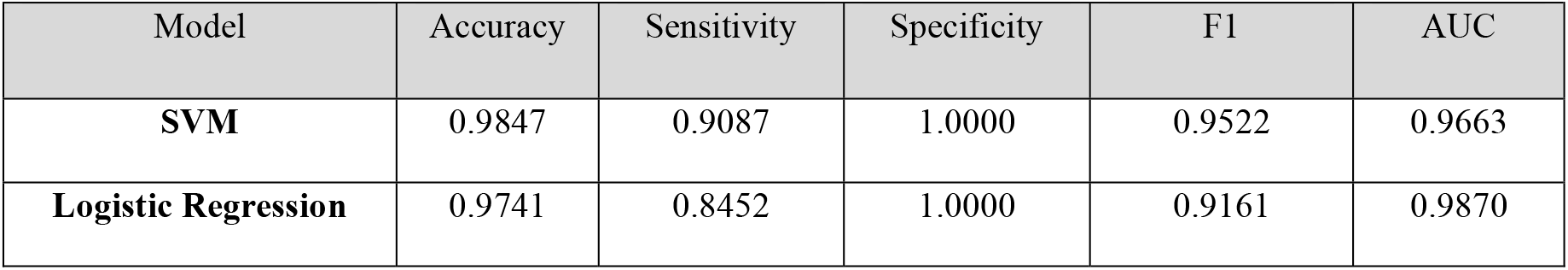
Performance of SVM model and LR model in classification and prediction of perceptual outcomes in individual trials of ‘attentional-blink’ task based on the features of pupil dynamics projected on two-dimensional principal component space.

Figure 3A shows fluctuations in the normalized pupil diameter averaged across blink (magenta) and no-blink (blue) trials, as illustrated in Figure 1D, smoothened by fitting a pair of gaussian functions mentioned above (fit parameters: *a1* = 1.02, *b1* = 1960.2, *c1* = 9482.4, *a2* = 0.021, *b2* = 153.09, *c2* = 330.45; *R^2^* = 0.969 for no-blink trials, and *a1* = 1.01, *b1* = 1933.6, *c1* = 12895, *a2* = 0.014, *b2* = 230.05, *c2* = 259.69; *R^2^* = 0.889 for blink trials). Using a hand-held pupillometer (https://www.idmed.fr/en/pupillometry/), we recorded pupil diameters of nine healthy young people who did not participate in the RSVP task. Figure 3B shows the variations in the average (± SEM) pupil diameter in millimetre over time. We subsequently derived a pupillary impulse response function (puIRF), 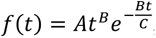, where the fit parameter C (= 16.67 ms) represented the time of occurrence of maximum impact on pupillary aperture in response to visual transients, and was set to the minimum duration of a flash of light on a 60 Hz monitor; *B* (= 1.0428) represented the recovery rate, which was derived from the best fit of the average pupil diameter shown in Figure 3B by an Erlang Gamma function [15,16]; and *A* (= – 0.01) was an arbitrary scale factor (Figure 3C). The duration of puIRF was set to 83 milliseconds, which was the SOA between two consecutive items in the RSVP.

**Figure 3:**
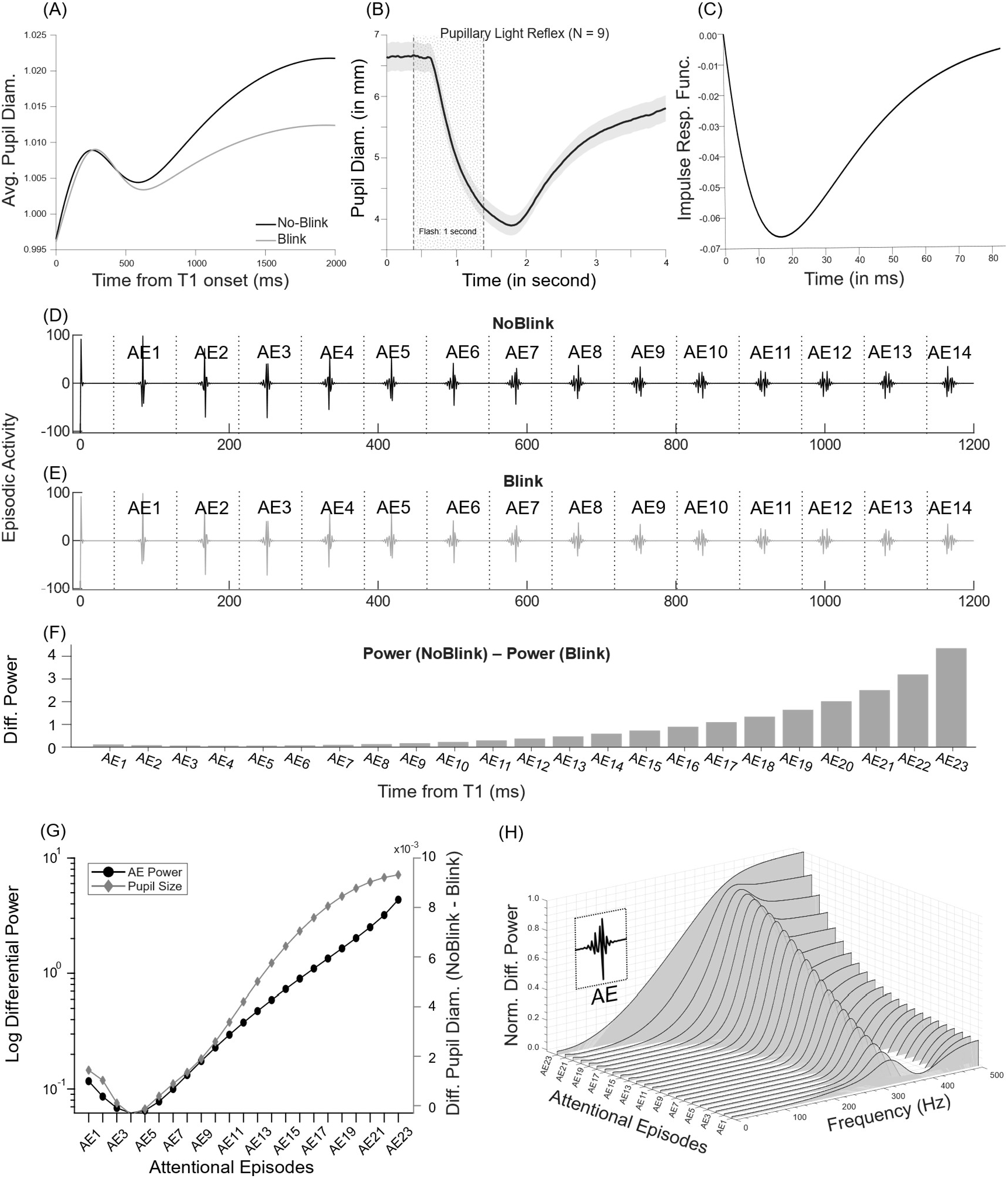
**(A)** Average normalized pupil responses in no-blink (black) and blink (grey) trials, each fitted by a pair of Gaussian function. (**B)** Average (± SEM) pupillary response elicited by a brief flash of light measured by a pupillometer. **(C)** Pupillary impulse response function (puIRF), modelled as an Erlang gamma function with the recovery rate obtained from the pupil-dynamics shown in (B). **(D), (E)** Attentional Episodes (AEs) reconstructed by deconvolving the gaussian-fitted pupil response in No-Blink and Blink condition, respectively, shown in (A) with the puIRF shown in (C). **(F)** Difference in root mean square (RMS) power between corresponding attentional episodes in no-blink and blink conditions [Power (No-blink) – Power (Blink)], increased progressively across successive episodes following T1 onset. **(G**) Logarithmic differential RMS power (black circles) and differential pupil diameter (grey diamonds) between no-blink and blink conditions exhibit highly correlated temporal dynamics. (H) Spectrogram of normalized differential AE power across frequencies for AEs. Power magnitude and occupied bandwidth (122–494 Hz) increased over time after T1 onset.

Figure 3D and E show signals obtained naturally from the average pupillary response in no-blink and blink trials shown in Figure 3A, respectively, after deconvolution by the puIRF shown in Figure 3C. We argue that the puIRF represented the intrinsic dynamics of the pupillary system in response to a brief flash of light and functioned as a temporal filter that transformed attentional episodes (AEs) caused by elements in the RSVP stream to the observed modulation in pupil size. Thus, the deconvolution technique reconstructed AEs from the fluctuations in the pupillary aperture in no-blink (Figure 3D) and blink (Figure 3E) condition. The root mean square (RMS) power of a signal, which is a measure of its effective magnitude or energy content over time (i.e., for a continuous signal *x(t),* over a period T, power 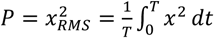 was calculated for each AE. We found that, AEs in no-blink trials contained more power than blink trials did in the corresponding AEs. Figure 3F shows that the difference between power in no-blink and blink trials steadily increased as time progressed after T1 onset. Interestingly, the dynamics of differential power in logarithmic scale resembled the difference in pupillary dynamics between no-blink and blink trials as shown in Figure 3G. Spearman’s Rank Correlation test indicated that dynamics of AE power and pupil size were highly synchronized (ρ = 0.998, r_s_^2^ = 0.996, p < 0.001, effect size = 0.07). The spectrogram in Figure 3I illustrates the distribution of differential power normalized to 1, across frequencies for different AEs. The magnitude and spread of power of the differential AE signal increased as the time progressed after T1. The ‘occupied bandwidth’ or the frequency bandwidth that contained 99% of a signal’s total integrated power was between 122 and 494 Hz.

## Discussion

In this study, we introduce a technique that can predict perceptual outcomes from pupillary dynamics in ‘attentional blink’ task on a trial-by-trial basis. To this end, we used machine learning techniques, specifically the ‘Support Vector Machine’ (SVM) and ‘Logistic Regression’ (LR), which classified blink and no-blink trials with high accuracy. We show that the differential power of attentional signals, reconstructed by deconvolving pupil dynamics, in the logarithmic scale is highly correlated with difference in pupil size between no-blink and blink trials. During the period when the attentional blink (AB) effect is most severe, i.e., between 200 and 400 ms after T1, the differential power of the signal is minimum and close to zero. As time progresses the differential power increases exponentially, thus the probability of detection of T2 also increases. This is consistent with the idea of ‘hazard rate’ in the context of temporal attention, which represents the conditional probability that the target will occur at a particular moment given that it has not yet occurred, and increases over time when the occurrence of the later target becomes progressively more likely [17,18].

In RSVP paradigms, the detection of T1 is thought to trigger an episode of attentional allocation that facilitates the selection and consolidation of T1 into capacity-limited working memory (WM) while simultaneously modulating the processing of subsequent stimuli [19]. According to episodic theories, the AB emerges because attentional control systems transiently suppress or gate incoming information to protect the integrity of the ongoing episode [20,21]. Consequently, stimuli arriving shortly after T1, including T2, may fail to achieve conscious access if they fall outside the optimal temporal boundaries of the episode. More recent computational models of the AB propose that attention operates in discrete “attentional episodes” (AE), delimited by specific events relevant for perception, rather than as a continuous process [12,22]. The AEs refer to transient windows of enhanced neural and cognitive engagement initiated by task-relevant events. The episodic simultaneous type/serial token (eSTST) model explains AB as a consequence of creating episodic representations (“tokens”) for targets in RSVP. According to the model, attentional enhancement triggered by T1 temporarily suppresses new AEs to preserve episodic distinctiveness, which can impair perception of T2 when it appears shortly afterward, whereas longer lags allow initiation of a new episode and successful WM encoding [10,23].

Neurophysiological evidence also suggests that selection in the WM and attention to sensory stimuli are mediated by shared neural mechanisms [24]. There is converging evidence that different parameters of alpha oscillations (8–12 Hz), which indicate internally and externally oriented brain states, may be critical for AB performance, as they can regulate attentional control and suppress distractor interference [25–27]. Note that usually RSVP elements also appear at a frequency within the same frequency range as the alpha oscillation. Contingent attentional capture theory suggests that attention is preferentially captured by goal-relevant distractors [28–31]. Additionally, no physiological evidence for active suppression of distractors or T2 after T1 processing during the RSVP stream is thus far available, which further supports the possibility of AEs corresponding to each item in the RSVP stream [32–34]. This episodic enhancement of attention is linked to activity of Locus Coeruleus (LC), which exhibits a distinct refractory period following phasic activity elicited by target stimuli [6]. The LC activity is tightly coupled with changes in the size of the pupillary aperture [5,35], and transient pupil dilations have consistently been linked to successful target detection across sensory modalities [36–38]. Larger pupil diameter preceding near-threshold targets has also been associated with improved perceptual accuracy under varying luminance conditions [39–41], while greater task-evoked pupillary responses correlate with enhanced behavioural performance [42] and reduced decision uncertainty [43,44]. Consistent with these findings, we observed larger pupil dilations during no-blink trials compared with blink trials, suggesting that successful conscious access to T2 is associated with enhanced attentional gain and task engagement.

A physiological account of AB, the Locus Coeruleus–Norepinephrine (LC-NE) model attributes the limitation of processing a stimulus when processing of another one is in progress, to the refractory dynamics of the LC. In response to T1, the LC exhibits phasic activation, transiently enhancing cortical processing through NE release. However, this release is autoinhibitory, producing a quiescent period of approximately 300 – 450 ms after T1 onset. Consequently, T2 presented within this interval undergoes weakened processing [9].

Attentional processes are also reflected in P300 or P3 EEG event-related potentials, particularly the parietal P3b component, which is modulated by phasic activity of the LC – NE system, and linked to attentional selection, stimulus categorization, and working-memory updating [45]. This model partially explains our findings, as it primarily emphasizes T1-related processing limitations. In contrast, our results showed no significant differences in T1-evoked pupillary responses between blink and no-blink trials. Instead, significant divergences emerged after T1 offset and during T2 processing. Thus, while our findings are broadly consistent with the ‘Episodic Distinctiveness’ and ‘LC–NE’ accounts of the AB phenomenon, they also suggest novel aspects that may be incorporated into existing models.

Given that the pupil dynamics did not show any major differences in T1 processing between blink and no-blink trials, and divergence emerged after the offset of dilation related to T1 processing, the eSTST framework provides a better interpretation of the data. Instead of viewing the attentional blink as a processing bottleneck caused by T1, the eSTST model argues that the blink is an adaptive mechanism that creates episodic distinctiveness in attention and working memory. In contrary to this model’s assumption that AEs are only triggered by targets and are relatively prolonged, our results indicate each item in RSVP stream is attentionally scanned, before selection of T2 (12). The occurrence of each AE is expressed in some kind of oscillatory activity, and their temporal dynamics appear to be tuned to the RSVP presentation rate. Note that, this shorter timescale of AEs is consistent with evidence from EEG studies highlighting the rhythmic nature of attentional selection [46].

Deconvolution analyses further revealed differential attentional adjustments to intervening distractors between ‘blink’ and ‘no-blink’ Condition, providing additional evidence for the contribution of distractor processing in AB. In RSVP task, attentional events occur rapidly, but the pupil responds slowly and smoothly. We derived an impulse response function (IRF) that describes how the iris muscle-plant responds over time to an unitary stimulation (i.e., brief flash of light) [47,48], which characterized the pupillary system’s intrinsic dynamics and acted like a temporal filter that transformed the original signal (i.e., attentional episodes) into the observed output (i.e., fluctuations in pupil size) [13,15,49,50]. In mathematics, the goal of deconvolution is to reconstruct the original signal as it existed prior to convolution by an IRF. We deconvolved pupillary response measured during the task performance by the IRF to retrieve the timing and magnitude of the underlying attentional events.

Neural responses to sensory information often depend more on relative than absolute changes. This compressive coding strategy, where information is represented on a logarithmic scale instead of a linear scale, is thought to enhance efficiency, sensitivity, and information storage in the nervous system [51]. Evidence for logarithmic computation by neurons has been observed in sensory perception [52], numerical cognition [53], time perception [54], and decision-making [55]. Here, we show that the logarithmically scaled difference in the episodic power of the oscillations obtained through deconvolution of pupil dynamics between no-blink and blink trials statistically closely corresponds to the linear difference in pupil dynamics between these trial types. This finding suggests that these oscillations may reflect neural encoding of attention involved in modulating pupillary aperture size. While previous studies employed temporal principal component analysis (PCA) [56–59], independent component analysis (ICA) [60], and related signal decomposition methods to reveal latent cognitive processes such as surprise, cognitive load, and working memory, this is the first study, to the best of our knowledge, that shows how perception may be predicted from pupillary dynamics as outcomes of modulations in temporal attention on single-trial basis.

## Conclusion

We addressed how pupillary dynamics can reliably predict decisions requiring supraliminal perception. Following correct identification of the first target when the second target was detected in a rapid stream of distractors, pupils dilated significantly more in comparison to when it escaped conscious perception due to temporal limitations of attention. Features extracted from progressive pupil-size fluctuations across trials in both scenarios were projected onto a lower-dimensional space via principal component analysis. Subsequently we classified and cross-validated those features using supervised machine-learning algorithms, achieving near-perfect prediction of perceptual decision outcomes. Furthermore, the deconvolution of average pupillary dynamics reconstructed a series of brief spells of oscillations mimicking attentional episodes associated with every item in the rapid stream of stimuli. We found that the relative strength of the attentional episodes was linked to the likelihood of the subsequent target to be perceived. These results show that pupil size is a potential marker of conscious perception on a trial-by-trial basis at finer time-scale.

## Data Availability Statement

The data and analysis code supporting the findings of this study will be made available to reviewers upon request in a format suitable for the peer-review process. Upon acceptance of the manuscript, all raw data and associated analysis code will be deposited in a publicly accessible repository (e.g., GitHub) and made freely available without restriction.

